# Porcine commensal *Escherichia coli*: A reservoir for class 1 integrons associated with IS*26*

**DOI:** 10.1101/158808

**Authors:** Cameron J. Reid, Ethan R. Wyrsch, Piklu Roy Chowdhury, Tiziana Zingali, Michael Liu, Aaron Darling, Toni A. Chapman, Steven P. Djordjevic

**Affiliations:** The ithree institute, University of Technology Sydney, Ultimo, NSW, 2007, Australia; NSW Department of Primary Industries, Elizabeth Macarthur Agricultural Institute, Menangle, NSW, 2568, Australia.

**Author notes:** These authors contributed equally to this work (Cameron J. Reid, Ethan R. Wyrsch). Corresponding author: Steven P. Djordjevic The ithree institute, University of Technology Sydney, PO Box 123, Broadway, Sydney, 2007.

## Abstract

Porcine faecal waste is a serious environmental pollutant. Carriage of antimicrobial resistance and virulence-associated genes (VAGs) and the zoonotic potential of commensal *Escherichia coli* from swine is largely unknown. Furthermore, little is known about the role of commensal *E. coli* as contributors to the mobilisation of antimicrobial resistance genes between food animals and the environment. Here, we report whole genome sequence analysis of 141 *E. coli* from the faeces of healthy pigs. Most strains belonged to phylogroups A and B1 and carried i) a class 1 integron; ii) VAGs linked with extraintestinal infection in humans; iii) antimicrobial resistance genes *bla_TEM_, aphAl, cmlA, strAB, tet(A)*A, *dfrA12, dfrA5, sul1, sul2, sul3*; iv) *IS26;* and v) heavy metal resistance genes (*merA, cusA, terA*). Carriage of the sulphonamide resistance gene *sul3* was notable in this study. The 141 strains belonged to 42 multilocus sequence types, but clonal complex 10 featured prominently. Structurally diverse class 1 integrons that were frequently associated with IS26 carried unique genetic features that were also identified in extraintestinal pathogenic *E. coli* (ExPEC) from humans. This study provides the first detailed genomic analysis and point of reference for commensal *E. coli* of porcine origin, facilitating tracking of specific lineages and the mobile resistance genes they carry.

**Conflict of Interest Statement:** None to declare.

## Introduction

*Escherichia coli* is the most frequently isolated Gram-negative pathogen affecting human health (Poolman and Wacker, 2016). They are frequently resistant to multiple antibiotics and modelling studies forecast that multi-drug resistant (MDR) *E. coli* infections will account for 30% of 10 million fatal MDR infections annually by 2050 (O’Neill, 2016). In addition to the pathogenic variants, commensal *E. coli* comprise an important component of the gut flora. Commensals colonise the gastrointestinal tract of warm-blooded mammals and reptiles from which they are shed into the open environment where they can persist and colonise other host species (Tenaillon et al., 2010, van Elsas et al., 2011). *E. coli* are shed into the environment in high numbers, for example each gram of faeces from commercially-reared pigs contains between 10^4^ – 10^8^ *E. coli* (Herrero-Fresno et al., 2015). In China alone, an estimated 0.618 billion to 1.29 billion metric tonnes of faeces are produced by swine each year (Han, 2014, Wang et al., 2006).

Pathogenic *E. coli* are broadly divided into intestinal pathogenic *E. coli* (IPEC) and extraintestinal pathogenic *E. coli* (ExPEC). ExPEC have a faecal origin, having persisted asymptomatically in the gut before opportunistically colonising extraintestinal sites where they cause a diverse range of diseases including urinary tract infections, pyelonephritis, wound infections, sepsis and meningitis (Johnson and Russo, 2005). ExPEC are thought to have foodborne reservoirs (Lyhs et al., 2012, Manges, 2016, Nordstrom et al., 2013, Vincent et al., 2010) and may enter the food chain via a number of sources (Ho et al., 2015, Johnson et al., 2005b, Pradel et al., 2008, Rees et al., 2015, Rizzo et al., 2013, Schroeder et al., 2003, Sola-Gines et al., 2015, Varela et al., 2015, Zurfluh et al., 2015). The zoonotic potential of commensal porcine *E. coli* as a source of ExPEC that cause disease in humans is unknown. ExPEC cannot be reliably detected in a diagnostic test as they are yet to be shown to possess a unique identifying feature relative to other strains of *E. coli* (Cordoni et al., 2016). Instead, as we aim to do here, whole genome sequencing can be used to discriminate strains indistinguishable by other methods and identify any genetic relationships between *E. coli* strains isolated from humans and from pigs.

The *E. coli* genome comprises both core and accessory genetic information. Horizontally acquired DNA, mediated by mobile genetic elements, plays an important role in the evolution of the species. Commensal or pathogenic bacteria may, in a single horizontal gene transfer event, acquire a mobile genetic element carrying multiple ARGs, virulence genes, and other genetic cargo that encode traits that offer a niche advantage (Kingsley et al., 2009, Levings et al., 2008, Levings et al., 2005, Levings et al., 2007, Liu et al., 2015, Moran et al., 2016, Roy Chowdhury et al., 2015, Venturini et al., 2010, Venturini et al., 2013). Furthermore, indirect selection pressure can, in the absence of antibiotic use, lead to the persistence of transferred genes. For example, heavy metals such as copper and zinc in feed formulations for food animals exert pressure on metal resistance genes that co-localise with ARGs (Baker-Austin et al., 2006, Li et al., 2017). The release of MDR commensal *E. coli* into the environment, such as when pig faeces are used as fertiliser, ensures the transfer and uptake of mobile elements into microbial communities in a manner that is poorly understood. Antibiotic resistance genes cluster on mobile genetic elements forming complex antibiotic resistance gene loci (CRL). Selection pressure afforded by any one of a number of antibiotics and heavy metals (zinc, cadmium, mercury) that contaminate fecal waste or those used used in food-producing and hospital environments is suffient to select for retention of the CRL (Ben et al., 2017, Zhou et al., 2017). Antimicrobial residues are present in the environment due to excretion from animals and humans (He et al., 2016, Nguyen Dang Giang et al., 2015, Wang et al., 2015, Alcock et al., 1999). Understanding of how antimicrobial resistance genes assemble on mobile genetic elements and the extent to which these then traffic through human, food animal and environmental reservoirs remains limited.

Class 1 integrons are a reliable proxy for the presence of multiple ARGs within bacteria in clinical and veterinary settings (Gillings et al., 2015). They are defective transposons that can capture ARG cassettes from the environmental resistome and express them via a promoter residing in the class 1 integrase gene. They are often mobilised by mercury resistance transposons belonging to the Tn2*1* family, which have been disseminated globally on a wide variety of conjugative plasmid backbones (Liebert et al., 1999). Resistance genes can also be acquired, lost and rearranged in bacteria by genetic events that involve insertion sequences (IS) such as IS*26,* IS*Ecpl* and IS*CR1* (Cheng et al., 2016, Harmer and Hall, 2016, Harmer et al., 2014, Toleman et al., 2006). IS26 is prominent in this regard because it has no target site specificity and can mobilise an increasing variety of ARGs (Canton and Coque, 2006, Dionisi et al., 2009, Dropa et al., 2016, Fournier et al., 2006, Harmer et al., 2014, Liu et al., 2017, Roy Chowdhury et al., 2015, Toleman and Walsh, 2011). Furthermore, IS26 is recognised to play a key role in: i) the evolution of plasmids and genomic islands that carry combinations of virulence and antimicrobial resistance genes (Doublet et al., 2009, Siebor and Neuwirth, 2013, Venturini et al., 2010, Venturini et al., 2013); ii) driving the formation of cointegrate plasmids encoding virulence and antibiotic resistance genes (Mangat et al., 2017) and; iii) initiating deletions in large multidrug resistance plasmids that enhance plasmid stability and expand host range (Porse et al., 2016).

Infectious disease management relies on surveillance of antimicrobial resistance and emerging pathogens using a one health approach. There is currently no published data available that records resistance gene or virulence gene carriage in commensal *E. coli* from Australian pigs. Here, for the first time, we present whole genome sequence analysis of 141 MDR commensal *E. coli* from pigs commercially-reared in Australia. We present data characterising their phylogenetic diversity, carriage of virulence genes, resistance genes and an analysis of the class 1 integrons they carry.

## Methods

### Management of farms and animals

The study was conducted using *E. coli* sourced separately from two pig production farm systems located approximately 250 km apart. Farms were designated descriptors F1 and F2. At both farms, pigs are intensively housed and kept in total confinement. Both farms have used neomycin in the past for the treatment of diarrhoeal disease. No antibiotics were being used during the first sampling time at F1, however the several pigs sampled at the second sampling time had received a course of neomycin (see below). No antibiotics had been administered to the pigs at F2 prior to sampling.

### Escherichia coli strains used in the study

*Escherichia coli* isolates were collected via rectal swab sampling using pigs between 19 and 30 days of age. At farm 1, rectal swabs were collected in May 2007 from pigs that were infected with enterotoxigenic *E. coli* (ETEC) during an outbreak of diarrhoeal disease, and then again in June 2007, when the disease had subsided after neomycin treatment and the next weaner batch was transferred into the weaner shed. At farm 2, rectal swabs were performed on healthy weaners that had not previously suffered diarrhoeal disease.

*E. coli* were isolated at the Elizabeth MacArthur Agricultural Institute (EMAI). Up to ten *E. coli* colonies were selected from individual pigs using MacConkey Agar. The total collection from farm 1 was 164 isolates whilst farm 2 was 171 isolates. All strains were screened by PCR for the class 1 integrase gene *intI1.* A subset of 141 strains that were positive for the *intI1* gene were included in this study. The subset of strains that were sequenced were identified as F1 + F2 (n = 141 from 50 pigs). This subset consisted of 94 strains from 21 pigs sampled at farm 2 (strains designated F2), and 47 strains from 29 pigs sampled at farm 1 (strains designated F1); among the F1 strains, 24 were disease associated (isolate numbers 1-30) and 23 were isolated from healthy animals (isolate numbers 365-409) Only 13 isolates in the collection carried toxin genes associated with porcine ETEC.

F1 strains were tested at EMAI using the calibrated dichotomous susceptibility test (CDS) for resistance to 12 antibiotics (Bell et al, 2013). The following were tested: ampicillin (25 μg), cefoxitin (30 μg), nalidixic acid (30 μg), ciprofloxacin (2.5 μg), imipenem (10 μg), sulphafurazole (300 μg), trimethoprim (5 μg), tetracycline (10 μg), neomycin (30 μg), gentamicin (10 μg), azithromycin (15 μg) and chloramphenicol (30 μg). This data is available in Supplementary Table 1 for strains sequenced in this study. F2 strains were tested for resistance to antibiotics at The ithree institute, University of Technology Sydney using the same method and panel of antibiotics as the F1 collection. F1 strains were also tested with streptomycin (25 μg) and kanamycin (50 μg). This data is available in Supplementary Table 1 for strains sequenced in the study. For storage, all strains were freshly cultured in LB-broth and frozen as glycerol stocks made using 500 μl of M9 salts solution and 500 μl of 50% glycerol and stored at -80°C. All strains were cultured in LB-broth prior to isolation of genomic DNA used for sequencing. Two enterotoxigenic *E. coli* (ETEC) strains, which were submitted to EMAI from Australian veterinary services as clinical, pig-derived strains were included in the phylogenetic analysis as reference strains.

### DNA extraction, whole genome sequencing and assembly

Genomic DNA was extracted using the ISOLATE II Genomic DNA Kit (Bioline, Eveleigh, Australia) following the manufacturers standard protocol for bacterial cells and stored at – 20°C. Whole genome sequencing libraries were prepared from separate aliquots of sample DNA using the Illumina Nextera DNA kit with modifications. In brief, the genomic DNA was first quantified using a Qubit ds DNA HS Assay Kit (Thermo Fisher Scientific, Scoresby, Australia). All sample gDNA concentrations were standardised to equal concentration to achieve uniform reaction efficiency in the tagementation step. Standard Illumina Nextera adaptors were used for sample tagmentation. The PCR-mediated adapter addition and library amplification was carried out using customized indexed i5 and i7 adaptor primers (IDT, Coralviulle, IA, USA), which were developed based on the standard Nextera XT Indexed i5 and i7 adapters (e.g. N701-N729 and S502-S522). Libraries were then pooled and size selected using SPRI-Select magnetic beads (Beckman Coulter, Lane Cove West, Australia). Finally, the pooled library was then quality checked and quantified on an Agilent Bioanalyzer 2100 using the DNA HS kit (Agilent, Santa Clara, CA, USA). Whole genome sequencing for the majority of F1 strains and ETEC strains was performed as previously reported (Darling et al., 2014b) using an Illumina MiSeq sequencer and MiSeq V3 chemistry. Whole genome sequencing of the remaining F1 and F2 strains was performed using an Illumina HiSeq 2500 v4 sequencer in rapid PE150 mode (Illumina, San Diego, CA, USA).

Sequence read quality was initially assessed using FastQC version 0.11.5 (http://www.bioinformatics.babraham.ac.uk/projects/fastqc/). Illumina raw reads passing quality control were assembled into draft genome sequences using the A5 assembly pipeline version A5-miseq 20140604 (Coil et al., 2015). Genome sequences have been deposited in the European Nucleotide Archive with study accession number PRJEB21464. Individual accession numbers for each sample are listed in Supplementary Table 1.

### Assembly Statistics

Comprehensive assembly statistics for the 143 porcine-derived *E. coli* are available in Supplementary Table 1. The number of scaffolds per genome ranged from 29 to 786, with a mean of 229. Each genome sequence had a median sequencing coverage of at least 20 × with a maximum of 94 × and mean of 54 ×.

### Gene identification and serotyping

Resistance, virulence and plasmid-associated genes were identified using in-house databases of virulence, resistance, plasmid replicon and OH-antigen genes and local BLASTn v2.2.30+ (Camacho et al., 2009) searches with an e-value of 1.0 × 10^−3^. Genes were considered present if the subject nucleotide sequence was > 90% identical over 100% of the length of the query sequence. BLAST hits covering less than 100% of the query were considered positive if they were truncated by a scaffold break or insertion. In-house databases were generated from GenBank sequences, as well as publicly available databases such as ResFinder, VirulenceFinder, ISfinder, SerotypeFinder (Zankari et al., 2012, Siguier et al., 2006, Joensen et al., 2015, Joensen et al., 2014), all accessible from the Centre for Genomic Epidemiology (CGE) (http://www.genomicepidemiology.org/).

### Phylogrouping and eMLST

*E. coli* phylogroups were determined using the scheme published by Clermont *et al,.* (Clermont et al., 2000). The genes *chuA* (gb|U67920.1), *yjaA* (gb|NC_000913.3) and the DNA fragment TspE4.C2 (gb|AF222188.1) were sourced from GenBank and identified *in silico* using BLASTn. Multilocus sequence typing (MLST) was performed *in silico* (eMLST) using the PubMLST database (http://pubmlst.org/) and the Achtman *E. coli* MLST scheme (http://mlst.warwick.ac.uk/mlst/).

### Phylogenetic Analyses

Maximum-likelihood phylogenetic distances between genomes were analysed using the PhyloSift pipeline (Darling et al., 2014a), and a tree was generated using FastTree2 (Price et al., 2010) and FigTree v1.4.2 (http://tree.bio.ed.ac.uk/software/figtree/). The FastTree2 protocol was modified to resolve short branches as described previously (Wyrsch et al., 2015)

## Results

Our study collection consisted of 141/316 (45%) strains of *E. coli* isolated from rectal swabs of pigs from two farms in New South Wales, Australia that were PCR-positive for the class 1 integron integrase gene, *intI1.* Initial screening indicated that 117/164 (71%) *E. coli* from farm 1 and 168/171 (98%) from farm 2 carried *intI1.* Such high carriage rates suggest that commensal *E. coli* from swine are resistant to multiple antimicrobial agents. Whole genome sequencing of 141 strains from F1 and F2 was employed to examine phylogeny and to determine the carriage of resistance and putative virulence genes among commensal *E. coli* sourced from these two commercial swine farms.

### Population structure of E. coli isolated from pig rectal swabs

We classified the strains in our study collection by phylogrouping, *in silico* MLST (eMLST), and *in silico* serotyping (e-serotyping). The majority of the strains in our study collection [103/141 (73%)] belonged to phylogroup A while the reminder belonged to phlylogroup B1 (21; 15%), phylogroup B2 (7; 5%), and phylogroup D (10; 7.1%). We identified 42 distinct sequence types, 21 of which were previously isolated from swine, as reported by the *E. coli* MLST database (http://mlst.warwick.ac.uk/mlst/dbs/Ecoli; accessed June 2017). Only seven sequence types were common to both F1 and F2 (Figure 1, Figure 2 and Supplementary Table 1). The most prominent sequence types were ST10, ST48 and ST218, all of which are members of clonal complex 10 (CC10), followed by ST361, ST542 and ST641. Twenty-six STs were represented by a single isolate. A designation of non-typable (NT) was given to the 17 strains for which one or more alleles could not be determined. *In silico* O:H typing identified 57 complete serotypes; 10 strains were O-non-typable (Figure 1, Figure 2 and Supplementary Table 1). In general, strains of any given ST carried the same O:H alleles, though intra-sequence type variability was observed among more abundant STs belonging to CC10 and ST542.

**Figure 1:**
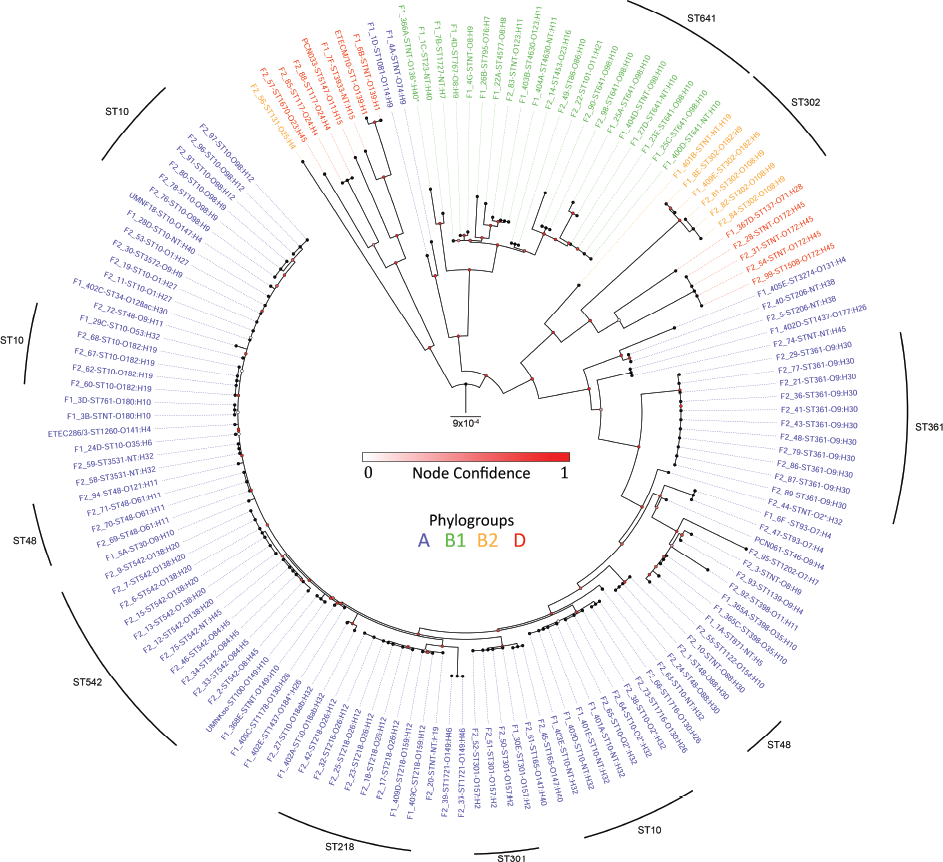
A mid-point rooted, maximum-likelihood phylogenetic tree inferred using PhyloSift v1.0.1, FastTree2 and FigTree v1.4.2. The tree contains all pig *E. coli* isolates sequenced in this study, plus four reference pig-sourced sequences. Tip colouring shows phylogrouping data (A-blue, B1-green, B2-orange or D-red) with names also containing MLST and serotype data. Node confidence values have been coloured on a 0 to 1, white to red scale. The tree scale (centre) shows the distance for 9 substitutions per 10 000 sites in the analysis. Dominant sequence types have been marked as an outer line.

**Figure 2:**
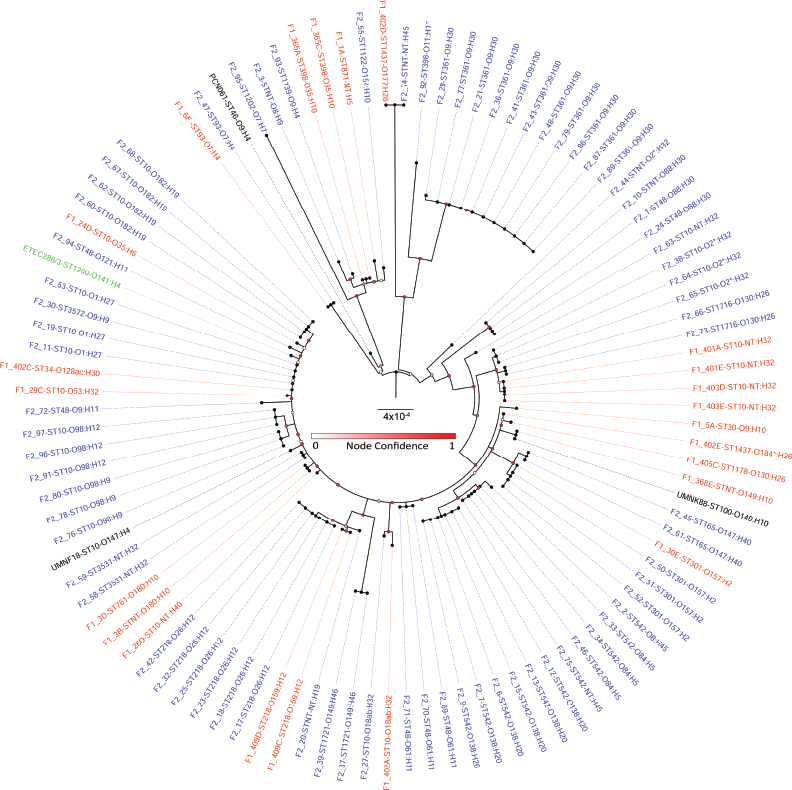
A mid-point rooted, maximum-likelihood phylogenetic tree generated from PhyloSift v1.0.1. The tree is composed of isolates from the phylogroup A clade that required higher resolution. Tips have been coloured according to source; F1 (blue), F2 (red), ETEC (green) and *E. coli* reference genomes (black). Node confidence values have been coloured on a 0 to 1, white to red scale. The tree scale (centre) shows the distance for 0.0004 substitutions per site. Outer line indicates clade groupings.

### Phylogenetic analysis

To determine genetic relatedness, we used PhyloSift, FastTree2 and FigTree v1.4.2 to generate and visualise a mid-point rooted, maximum-likelihood phylogenetic tree containing the F1 + F2 pig *E. coli* draft whole genome sequences, two ETEC strains and 4 publicly-available, pig-pathogenic *E. coli* complete genome sequences: *E. coli* UMNK88 (NC_017641.1), UMNF18 (NZ_AGTD01000001.1), PCN033 (NZ_CP006632.1) and PCN061 (NZ_CP006636.1) (Figure 1). Tree topology was highly congruent with Achtman MLST and *in silico* serotyping, grouping strains by sequence type, and then further by serotype. Clade structure was generally congruent with phylogroup analyses; however, eight strains belonging to phylogroups B2 and D formed a separate clade. We identified three major clades, with the eight B2/D phylogroup strains forming one of them. Of the other two clades, the first consisted almost exclusively of phylogroup B1 strains, with ST641 being the dominant sequence type, although two phylogroup A isolates (F1_1D and F1_4A) were outliers. The second was composed of two separate sub-clades, one with B2 and D strains (ST302 and ST1508:O172:H45 strains dominated), and the other with phylogroup A strains (ST10 and sequence types within CC10, as well as ST361 and ST542, isolates that were common in our study collection). To explore the diversity of the dominant phylogroup A strains an identical Phylosift analysis was performed on these strains. This tree revealed divergence among members of CC10 and provided higher resolution of varying serotypes within identical sequence types (Figure 2).

Branch distances within the mid-point rooted alignment indicated that B2 and D strains are evolutionarily distant from the dominant phylogroup A strains. Despite this, Figure 2 also demonstrates variability within phylogroup A and CC10, albeit with low confidence scores at some of the major split points.

### Distribution of antimicrobial resistance genes (ARGs) and heavy metal resistance genes

From BLAST analyses, we observed up to 15 ARGs per strain in our collection. Only one strain (F2_76) produced no matches to any reference ARG although this isolate was resistant to streptomycin and neomycin. Notably, 88/141 strains (77%) carried eight or more ARGs. Surprisingly, strains belonging to phylogroup A carried the highest average number of ARGs (10 per strain). Strains belonging to phylogroup B1, B2 and D carried an average 7.6, 9.4 and 8.1 ARGs per strain, respectively. The most common ARGs among the strains in our collection were the penicillin resistance gene, bla_TEM-1_, (112/141; 79%); *aphA1,* encoding resistance to kanamycin and neomycin (97/141; 68%), the co-linked streptomycin resistance genes, *strA* and *strB* (96/141; 68%), and the tetracycline resistance gene *tetA* (95/141; 67%). Quinolone resistance genes *oqxAB,* which typically localize on plasmids (34/141; 24%) were less frequently identified. Genes encoding extended spectrum β-lactamases (ESBL), extended spectrum carbapenemases (ESC) and resistance to macrolides were not detected. Heavy metal resistance genes including the copper resistance gene *cusA* (141/141; 100%), the Tn*21* mercury resistance gene *merA* (86/141; 61%) and the tellurite resistance gene *terA* (56/141; 40%) were identified frequently.

ARGs were located in integron-associated gene cassettes for many of our strains. Cassettes carried by the majority of strains included aminoglycoside resistance genes, *aadA1* (90/141; 64%) and *aadA2* (91/141; 65%); the chloramphenicol resistance gene cassette, *cmlA,* (77/141; 55%) and the trimethoprim resistance gene cassette, *dfrA12,* 78/141 (55%). The trimethoprim resistance gene cassette *dfrA5* was also observed frequently 64/141 (45%). We identified sulphonamide resistance genes frequently, of which *sul11* is a feature of the 3'conserved segment (CS) region of typical clinical class 1 integrons. Notably, *sul3* was identified in more strains (81/141; 57%) than was *sul1* (65/141; 46%) or *sul2* (57/141; 40%). This data is summarized as a heat map in Figure 3.

**Figure 3:**
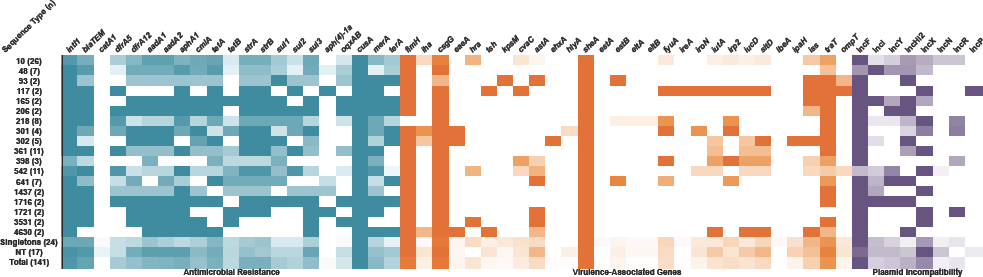
Heat map depicting carriage of antimicrobial resistance genes (aqua), virulence-associated genes (orange) and plasmid incompatibility groups (purple) by sequence type. Darker colour indicates high carriage amongst a given sequence type, lighter colour indicates lower carriage and white indicates no carriage. For full screening data see Supplementary Table 1.

### Multidrug resistant porcine E. coli carry structurally-diverse class 1 integrons

BLASTn analysis confirmed that 129/141 (91%) of F1 + F2 strains carried *intI1.* Notably, all 141 strains carried IS26. Among our study collection, we sought to identify regions featuring mobile elements in association with resistance genes. It is challenging to assemble complete sequences for such regions using Illumina sequence data because of the presence of repeated elements; however, we were able to determine the context of some of the most frequently occurring resistance genes in our collection. Numerous, structurally diverse class 1 integrons with altered *3‵-CS* associated with IS*26* were identified (Figure 4). We derived partial structures to most of the class 1 integrons, and IS26 flanked most of them. Notably, one of the structures (4C in Figure 4) was also identified in clinical *E. coli* strains from a Sydney hospital (data not shown).

**Figure 4:**
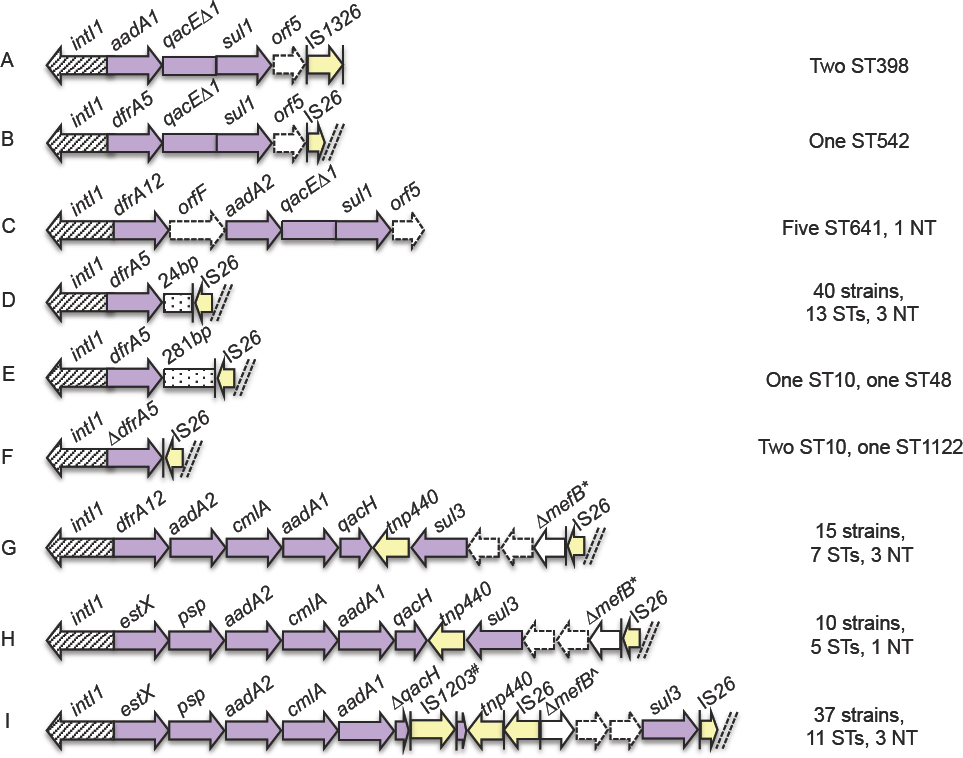
Schematic diagram (not to scale) of integrons within porcine strains that were sequenced. Arrows represent ORFs. Arrows with broken lines indicate hypothetical proteins. Vertical bars represent inverted repeats. Dashed double diagonal line represent sequence scaffold breaks. Antimicrobial resistance genes (purple) and IS elements (yellow), are colour coded. *indicates 260 bp of *mefB* remaining. ^A^ indicates 111 bp of *mefB* remaining. #IS1203-like

Three different class 1 integrons carried a *sul3* gene (4G, 4H and 4I in Figure 4). The first time that *sul3* was linked with *E. coli* from a food-animal source in Australia occurred in 2015 in a highly virulent porcine ST4245 ExPEC strain (Wyrsch et al., 2015). Moreover, *sul3* was first reported in a human in Australia in 2017 in a commensal *E. coli* with ST95 (Moran et al., 2017). In 4G and 4H, the *sul3* module, which comprises a putative transposase *tnp440*, *síti3,* two hypothetical proteins and 260 bp of the macrolide efflux gene *mefB* truncated by IS26, was the same. The 4G and 4H integron structures differed from each other in the cassette array upstream of the *sul3* module. The integron structure of 41 differed from that of 4G and 4H both in its *sul3* module, which carried an additional copy of IS*26*, length of the *mefB* gene fragment (111 bp) and an insertion of an IS*1203*-like element in *qacH.* Only three of the integrons among our strains (4A-C) carried a *sull* gene. These data suggest that Australian swine are a reservoir clinical class 1 integrons that carry *sul3* and have *3″-CS* that have been modified by IS*26*. Further studies of commensal and pathogenic strains of *E. coli* are needed to explore this hypothesis.

### Distribution of virulence-associated genes (VAGs) among pig faecal E. coli

To assess the virulence potential of commensal pig *E. coli* strains in our collection, we screened for a total of 67 genes that have been associated with either intestinal disease or extraintestinal disease caused by *E. coli* pathotypes. 29 of these genes were present in at least one isolate (See Figure 3 and Supplementary Table 1). All isolates possessed VAGs (range of 4 to 17 genes per strain), with the majority (98/141; 70 %) carrying between 4 and 8. The VAGs were present in diverse gene combinations between and within sequence types. From Phylogroup A, ST10, ST301, ST398, ST542, ST361 and ST1081 frequently carried more than five VAGs. Most VAGs were typical of extraintestinal *E. coli* pathotypes (ExPEC), whilst ETEC toxin gene carriage was only observed in 13 isolates.

### Plasmid incompatibility groups in porcine faecal E. coli

We screened the collection for plasmid replication-associated genes from nine plasmid incompatibility groups that are commonly associated with carriage and mobility of ARGs (Figure 3 and Supplementary Table 1). IncF was the most common replicon (114/141; 81%) followed by IncX (78; 55%) and IncHI2 (54; 38%). All replicons were present across multiple sequence types.

## Discussion

Globally, there is a poor representation of genomic sequences for commensal *E. coli* isolated from faecal populations of healthy pigs, and none in Australia. Here, for the first time, we sequenced the genomes of *E. coli* isolated from the faeces of healthy pigs and determined their Clermont phylogroup, MLST (Achtman), e-serotype as well as carriage of genes associated with virulence. The phylogenetic relationship shared by the 141 strains, the types of resistance genes that reside within the class 1 integrons, and how the structures of class 1 integrons are modified by IS*26* were also investigated. Despite sampling only two commercial piggeries, we identified a wide variety of multilocus sequence types. Our findings support the hypothesis that, even without direct selection pressure, commensal *E. coli* populations residing within the faeces of pigs are often resistant to multiple antimicrobial agents and carry numerous VAGs. Notably, we also identified genetic epidemiological markers for tracking CRL residing on mobile genetic elements in commensal *E. coli.*

### Commensal E. coli lineages are associated with disease

The dominant lineages in our collection, as determined by PhyloSift analysis, were phylogroup A *E. coli* belonging to sequence types residing within CC10, particularly ST10, ST48, ST218. Together with the observations of others, our data suggest *E. coli* of CC10 sequence type may be opportunistic, MDR pathogens with a broad animal host range. *E. coli* CC10 can colonise humans, swine, poultry, dogs, migratory birds, rodents, camels and cattle (Alcala et al., 2016, Baez et al., 2015, Cordoni et al., 2016, Ho et al., 2015, Liu et al., 2016, Manges, 2016, Manges et al., 2015, Rodrigues et al., 2013, Shabana et al., 2013, Trobos et al., 2009, Hasan et al., 2014). *E. coli* CC10 can also be isolated from raw and treated wastewater, and from urban streams (Varela et al., 2015). *E. coli* CC10 is increasingly associated with intestinal disease in humans (Guiral et al., 2011, Reuland et al., 2013) and extraintestinal infections in pigs (Ding et al., 2012, Tan et al., 2012), dogs (Liu et al., 2016), and humans, including UTI, pyelonephritis and sepsis (Giufre et al., 2012, Manges, 2016, Salvador et al., 2012, Usein et al., 2016). *E. coli* CC10 are often MDR, and the resistance genes they carry can encode resistance to extended-spectrum beta-lactams (Pires et al., 2016, Tagg et al., 2015). ST10 is a noted ExPEC sequence type in humans and has been identified in food animals, retail meats, and the environment (Alghoribi et al., 2015, Manges et al., 2015, Toval et al., 2014, Leverstein-van Hall et al., 2011, Moran et al., 2015). ST10 was the dominant sequence type in studies of porcine commensal *E. coli* samples in Denmark, Ireland and Spain (Cortes et al., 2010, Herrero-Fresno et al., 2015, Wang et al., 2016). The core attributes of ST10 that enable it to colonise diverse niches remain unknown. The phylogenetic diversity we observed within porcine faecal ST10 suggests that such attributes may vary between strains. Whole genome sequence analysis of *E. coli* ST10 genomes from different regions of the world and from different hosts is needed to understand the full diversity and success of this sequence type.

### MDR porcine E. coli carry atypical class 1 integrons

Notably, *sul3* was the most frequently identified *sul* gene in our collection and three different sul3-containing integron structures were identified. Carriage of class 1 integrons possessing *sul3* has been observed in disease-associated and commensal *E. coli* isolates from animals and humans, as well as in bacterial species other than *E. coli* from different countries (Antunes et al., 2007, Curiao et al., 2011, Liu et al., 2009). In Australia, the carriage of *sul3* by *E. coli* has been reported infrequently, although it has been identified in several uropathogenic *E. coli* isolates (Gundogdu et al., 2011), in a highly virulent porcine ST4245 ExPEC strain (Wyrsch et al., 2015), and in a human commensal ST95 *E. coli* on a virulence plasmid that carries multiple antibiotic resistance and virulence genes (Moran et al., 2016). In Europe, class 1 integrons containing *sul3* have been observed in commensal *E. coli* from both humans and animals, indicating they are widely disseminated in a variety of *E. coli* lineages (Bischoff et al., 2005, Partridge et al., 2009, Saenz et al., 2010, Sunde et al., 2008, Moran et al., 2016). Structures similar to ours have also been reported in *Salmonella enterica* serovars, suggesting inter-species transfer of class 1 integrons carrying *sul3* has occurred (Antunes et al., 2007). The majority of *sul3*-containing strains in our study collection carried *merA* derived from Tn*21* 48/81 (59%). This suggests that both Tn*21* and other elements may play a role in disseminating *sul3* integrons (Moran et al., 2016).

The potential role for *sul3*-integrons in intra-and interspecies exchange of antibiotic resistance makes it desirable both to understand their evolution and to track their movement through bacterial populations. In Figure 5, we provided a model that could explain the micro-evolutionary events that created the novel sul3-integron depicted in structure 4I. The 4I structure likely evolved from a progenitor similar to one described by Curiao *et al,* (GenBank entry gb|HQ875016.1), as this is the only report to describe *IS26* adjacent to *sul3* (Curiao et al., 2011). Conceivably, the novel 4I structure emerged from insertion of a second copy of IS26, which further truncated *mefB,* followed by an inversion event. To our knowledge, this is the first study to identify a 111-bp *mefB* variant. The 4I structure was observed within the collection in 37 *E. coli* strains of different sequence types, suggesting horizontal transfer of a mobile element(s) carrying the integron. Further work is needed to examine this hypothesis. IS*26*-mediated deletions of the *mefB* can be used to track *sul3*-containing integrons between *E. coli* populations. Firstly, one of a number of different truncated variants of the *mefB* gene is carried in most *sul3*-integrons found in human and animal derived *E. coli* (Antunes et al., 2007, Curiao et al., 2011, Liu et al., 2009, Moran et al., 2016). Secondly, the class 1 integrase upstream of the *sul3* module is likely to be functional based on our observation of different antibiotic cassette arrays associated with a 260-bp *mefB* deletion (see Figure 4G & 4H) in our study collection. BLASTn analysis identified *sul3* integrons carrying *ΔmefB* with a 260-bp deletion that are identical to those observed in porcine isolates P328.10.99.C2 and P528.10.99.C4 from Great Britain (Liu et al., 2009). Furthermore, plasmid pCAZ590 isolated from poultry in Germany carried an identical integron *(estX-psp-aadA2-cmlA-aadA1-qacI-tnp440-sul3-orf1-orf2-ΔmefB:260bp-IS26)* to 4H with an additional *bla_SHV-12_* gene 73 bp downstream of IS*26* (Alonso et al., 2017). Although the evolutionary events that lead to this derivative structure are not known, this plasmid illustrates how IS*26* augmented integrons may act as scaffolds for the integration of genes that confer resistance to critically important human antibiotics.

**Figure 5:**
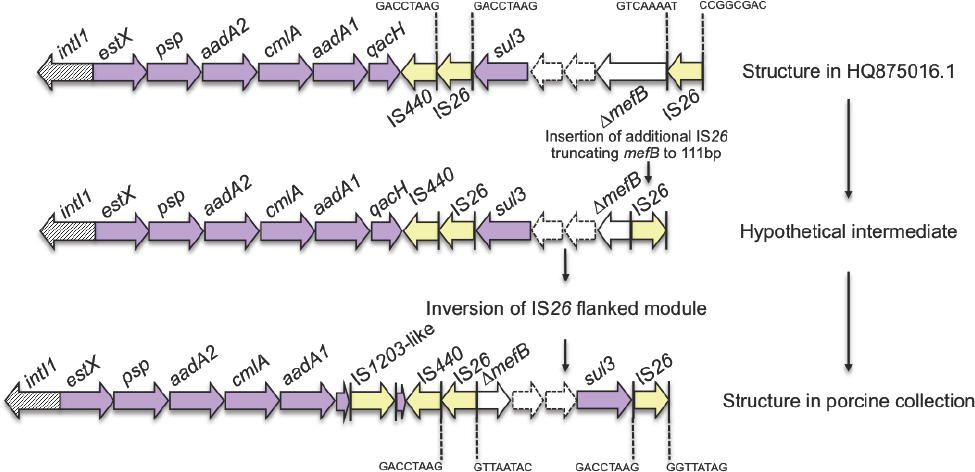
Schematic diagram (not to scale) of proposed evolutionary pathway to the *sul3-ΔmefB* arrangement shown in Figure 4I. IS*26* 8 bp direct repeats are annotated.

The deletion event in the 3″-*CS* of the integron depicted in 4E *(dfrA5-IS26)* may serve as another genetic signature for tracking resistance genes, and bacteria that carry them, through different hosts and environments (Djordjevic et al., 2013, Dawes et al., 2010, Levings et al., 2005, Roy Chowdhury et al., 2015). Previously we observed the 4E integron structure on plasmids carrying virulence genes in atypical EPEC strains isolated from cattle with gastrointestinal disease and *E. coli* strains linked to EHEC O26:H-isolated from humans with haemorrhagic colitis (Venturini et al., 2010, Venturini et al., 2013). In each of these earlier cases, the IS*26* that interrupted the 3″-*CS* of the integron formed part of the left boundary of Tn6026, an IS26-flanked, globally disseminated transposon that harbours multiple ARGs (Cain et al., 2010, Roy Chowdhury et al., 2015, Venturini et al., 2010, Venturini et al., 2013, Reid et al., 2015). Further studies are needed to determine if *Tn6029* or related transposon *Tn6026* are linked with the 4E structure in different *E. coli* in our collection.

### Zoonotic potential of commensal. E. coli from swine

In considering the zoonotic potential of pig faecal *E. coli,* we wanted to determine whether the strains in our collection carried VAGs that have been associated with the pathogenicity of *E. coli* linked to either intestinal or extraintestinal disease and, if those genes were in our strains, what proportion of the *E. coli* carry them? A limitation of investigating zoonotic potential for extraintestinal disease is the genetic redundancy identified in the virulence attributes from ExPEC. A recent study suggested that the number of virulence factors carried by an ExPEC strain is the only independent factor that can explain extraintesinal virulence in a mouse model of sepsis (Bleibtreu et al., 2013). Our collection contained strains possessing large numbers of VAGs, most notably B2-ST131 (n = 1) and D-ST117 (serotype O24:H4; n = 2). These two sequence types are representative of pandemic ExPEC clones that cause hospital and community-acquired infections in humans worldwide (Johnson et al., 2013, Stoesser et al., 2016, Manges et al., 2015). They have both been linked with poultry and have only rarely been isolated from porcine sources (Manges, 2016, Manges et al., 2015). The single ST131 strain in our porcine collection carried 10 antimicrobial resistance genes and 16 VAGs. The two ST117 strains carried eight antimicrobial resistance genes and 16 VAGs, including the full array of iron acquisition genes *fyuA, irp2, ireA, iroN, iutA, iucD* and *sitD.* Several of these genes are typically encoded on virulence plasmids circulating in APEC (Zhu Ge et al., 2014) and this profile is similar to O111:H4-ST117 strains from poultry previously reported by Mora *et al,* (Mora et al., 2012). The presence of ST117 and ST131 in our collection is intriguing and warrants further investigation.

Among other strains in our collection, we also identified genes that are associated with the ability to cause extraintestinal disease in humans as well as intestinal persistence (Johnson and Russo, 2005, Schierack et al., 2008), including genes that are under positive selection in uropathogenic *E. coli* (UPEC) (Chen et al., 2006), such as heat-stable agglutinin gene *hra* (Srinivasan et al., 2003), murine uroepithelial cell adhesin gene *iha* (Johnson et al., 2005a), iron acquisition genes, *fyuA, iutA, iucD* and *sitD;* and the serum survival genes *iss* and *traT.* Strains carrying *ibeA,* which has been linked with the ability of *E. coli* to invade brain microvascular endothelial cells and haematogenous meningitis (Che et al., 2011, Huang et al., 2001) and the intimin gene eaeA, that defines several intestinal *E. coli* pathotypes, including EHEC, EPEC and atypical EPEC (Kaper et al., 2004) were also identified.

The high frequency of VAGs in phylogroups A and B1, an average of 6.9 and 9.5 VAGs per isolate respectively, was unexpected because *E. coli* belonging to phylogroups A and B1 are considered to have low virulence potential (Ewers et al., 2007, Johnson and Stell, 2000). The carriage of multiple VAGs in pig *E. coli* is consistent with earlier studies (Dixit et al., 2004, Schierack et al., 2007). In China, ExPEC have been isolated from a variety of tissues and bodily fluids of pigs with septicaemia, meningitis and respiratory disease with increasing frequency since 2004 (Ding et al., 2012, Tan et al., 2012). It is notable that 35% of 81 isolates in the study of Ding et al., belonged to phylogroup A, clonal complex 10. In European wild boars, which are assumed to be ancestors of domestic pigs in Europe (Larson et al., 2005), *E. coli* strains carry, on average, seven or more VAGs, with some strains carrying up to 16 VAGs (Romer et al., 2012). Collectively, these observations suggest that *E. coli* phylogroup A and B1, at least those sourced from swine, carry multiple VAGs.

### Contribution of food production animals to the evolution of pathogens and antimicrobial resistance

MDR *E. coli* carrying ARGs associated with mobile genetic elements and VAGs are released into the environment by food production animals via release of faecal effluent. In Australia the capacity for pig production to contribute to the evolution and dissemination of pathogens and ARGs is more restricted compared to that of pig production systems in many other countries, due to a range of factors. Firstly, Australia has a large landmass that is surrounded by ocean preventing the movement of animals from neighbouring countries. Secondly, importation of food animals into Australia has been restricted since the 1970s (Turner, 2011). Thirdly, antibiotics such as fluoroquinolones cannot legally be administered to food animals and many others are restricted from use in food animal production (Cheng et al., 2012, Jordan et al., 2009). However, even in the restricted environment in Australia, phenotypic resistance to clinically-important antibiotics, including extended-spectrum cephalosporins and fluoroquinolones has been observed in *E. coli* which belong to globally disseminated *E. coli* lineages ST744, ST100 and ST1 (Abraham et al., 2015). Globally, genomic surveillance is needed to understand the relative contribution of food production animals to the complex web of interactions between microbiota and the mobile resistome; to provide baseline carriage rates for antimicrobial genes and VAGs; and to monitor the emergence of novel drug resistant pathogens (Wyrsch et al., 2016, Djordjevic et al., 2013).

In summary, we report the first genomic study of commensal *E. coli* isolated from commercial pigs used for food consumption and provide data to inform assessment of potential risks pig commensal *E. coli* may pose to human health. Such risks are complex and difficult to quantify. For example, attempts to attribute ExPEC transmission to humans from a food animal or environmental reservoir is complicated due to the delay between acquisition of an ExPEC pathogen in the human gastrointestinal tract and precipitation of a disease event (Manges, 2016). Nonetheless, it is well established that bacteria and the resistance and virulence genes they carry transfer between diverse animal hosts and environmental niches (Woolhouse et al., 2015). *E. coli* that evolve under antimicrobial selection pressures experienced in food production facilities persist on mobile elements that inhabit enterobacterial populations in humans (Anssour et al., 2016). Our results show that swine are a reservoir; i) for phylogroup A and B1 *E. coli* that carry VAGs, ii) the *sul3* gene; iii) atypical class 1 integrons associated with IS26; and iv) *E. coli* lineages belonging to CC10. Although strains in our collection did not carry resistance genes that render antibiotics critical to human health ineffective, they carry class 1 integron structures flanked by IS26. IS*26* is associated with mobile elements that carry many antimicrobial resistance genes, including those used to treat serious human infections, and it plays an important role in the assembly of resistance gene regions. Our study has identified several new genetic signatures that may be used in tracking mobile antibiotic resistance genes.

## Acknowledgments

This work is a product of the AUSGEM partnership. The work was supported in part by the Australian Government through the Australian Research Council, Linkage grant LP150100912. C.J.R and E.R.W are both recipients of Australian Government Research Training Program Scholarships. We thank Fiona MacIver for assistance with preparing the manuscript.

## Conflict of Interest

None to declare.

